# Targeted mutation detection in breast cancer using MammaSeq™

**DOI:** 10.1101/264267

**Authors:** Nicholas G. Smith, Rekha Gyanchandani, Grzegorz Gurda, Peter C. Lucas, Ryan J. Hartmaier, Adam M. Brufsky, Shannon Puhalla, Amir Bahreini, Karthik Kota, Abigail I. Wald, Yuri E. Nikiforov, Marina N. Nikiforova, Steffi Oesterrich, Adrian V. Lee

## Abstract

**Background:** Breast cancer is the most common invasive cancer among women worldwide. Next-generation sequencing (NGS) has revolutionized the study of cancer across research labs around the globe, however genomic testing in clinical settings remain limited. Advances in sequencing reliability, pipeline analysis, accumulation of relevant data, and the reduction of costs are rapidly increasing the feasibility of NGS-based clinical decision making.

**Methods:** We report the development of MammaSeq, a breast cancer specific NGS panel, targeting 79 genes and 1369 mutations, optimized for use in primary and metastatic breast cancer. To validate the panel, 46 solid tumor and 14 plasma circulating-free cfDNA samples were sequenced to a mean depth of 2311X and 1820 X respectively. Variants were called using Ion Torrent Suite 4.0 and annotated with cravat CHASM. CNVKit was used to call copy number variants in the solid tumor cohort. The oncoKB Precision Oncology Database was used to identify clinically actionable variants. ddPCR was used to validate select cfDNA mutations.

**Results:** In cohorts of 46 solid tumors and 14 cfDNA samples from patients with advanced breast cancer we identified 592 and 43 protein coding mutations. Mutations per sample in the solid tumor cohort ranged from 1 to 128 (median 3) and the cfDNA cohort ranged from 0 to 26 (median 2.5). Copy number analysis in the solid tumor cohort identified 46 amplifications and 35 deletions. We identified 26 clinically actionable variants (levels 1-3) annotated by OncoKB, distributed across 20 out of 46 cases (40%), in the solid tumor cohort. Allele frequencies of ESR1 and FOXA1 mutations correlated with CA.27.29 levels in patient matched blood draws.

**Conclusions:** In solid tumors biopsies and cfDNA, MammaSeq detects clinicaly actionable mutations (oncoKB levels 1-3) in 22/46 (48%) solid tumors and in 4/14 (29%) of cfDNA samples. MammaSeq is a targeted panel suitable for clinically actionable mutation detection in breast cancer.

## Background

Advanced breast cancer is currently incurable. Selection of systematic therapies is primarily based on clinical and histological features and molecular subtype, as defined by clinical assays [1]. Large-scale genomic studies have shed light into the heterogeneity of breast cancer and its evolution to advanced disease [2, 3], and coupled with the rapid advancement of targeted therapies, highlights the need for more sophisticated diagnostics in cancer management [4].

Next-generation sequencing (NGS) based diagnostics allow clinicians to identify specific putative driver events in individual tumors. Correctly identifying disease drivers may enable clinicians to better predict treatment responses, and significantly improve patient care [5]. However, to date, the use of NGS as a clinical diagnostic remains limited [6]. Published data regarding prognostic utility, and utilization for selection of targeted therapies or enrolment clinical trials is lacking.

The original 46 gene AmpliSeq Cancer Hotspot Panel (Thermo Fisher Scientific) was shown to have a diagnostic suitability in primary lung, colon, and pancreatic cancers [7], however, our previous report that surveyed the clinical usefulness of the 50 gene AmpliSeq Cancer Hotspot Panel V2 in breast cancer found that the panel lacks numerous key known drivers of advanced breast cancer [8]. For example, the panel does not include any amplicons in *ESR1*, which harbor mutations which are known to contribute to hormone therapy resistance (for review see [9]), and lacks coverage of the majority of known driver mutations in *ERBB2* [10].

The lack of any reported breast cancer specific diagnostic NGS test inspired the development of MammaSeq™, an amplicon based NGS panel built specifically for use in advanced breast cancer. 46 solid tumor samples from women with advanced breast cancer, plus an additional 14 samples of circulating-free DNA (cfDNA) from patients with metastatic breast cancer were used in this pilot study to define the clinical utility of the panel. The patient cohort encompassed all 3 major molecular subtypes of breast cancer (luminal, ERBB2 positive and triple negative), and both lobular and ductal carcinomas (Table 1).

**Table 1:** Patient and Specimen Characteristics.

|  | **Patients with available**<br>**tumor tissue (n=46)** |
| --- | --- |
| Age |  |
| Median age (yrs) | 45 |
| Range (yrs) | 31-71 |
| Race |  |
| White | 45 (97.8%) |
| Black | 1 (2.2%) |
| Site |  |
| Primary | 10 (21.7%) |
| Metastatic | 36 (78.3%) |
| Stage (Dx) |  |
| I | 10 (21.7%) |
| II | 8 (17.4%) |
| III | 13 (28.3%) |
| IV | 4 (8.7%) |
| Unknown | 11 (23.9%) |
| Hormone-receptor |  |
| HR + and HER2 – | 19 (41.3%) |
| HR + and HER2 + | 5 (10.9%) |
| HR + and HER2 Unknown | 1 (2.2%) |
| HR – and HER2 + | 1 (2.2%) |
| HR – and HER2 – | 17 (36.9%) |
| Both Unknown | 2 (4.3%) |
| Histopathology |  |
| Ductal | 34 (73.9%) |
| Lobular | 5 (10.9%) |
| Mixed | 3 (6.5%) |
| Other/Unknown | 4 (8.7%) |

## Methods

### Patient Sample Collection

For MammaSeq NGS testing, this study utilized breast tumors from 46 patients and blood samples from 7 patients. The research was performed under the University of Pittsburgh IRB approved protocol PRO16030066. The general patient characteristics are shown in Table 1 and more detailed patient information is shown in Supplemental Table 1. We utilized 46 of the 48 breast cancer cases previously described in a report by Gurda et al [8]. All of these cases underwent AmpliSeq Cancer Hotspot Panel v2 NGS testing between January 1, 2013 and March 31, 2015 within the UPMC health system. MammaSeq™ was performed on the identical genomic DNA isolated from these tumor specimens that was originally used for initial cinical testing. 2 cases were excluded due to insufficient DNA. In addition, a cohort of 7 patients with metastatic breast cancer (MBC) had 20ml venous blood drawn in Streck Cell-Free DNA tubes between July 1, 2014 and March 29, 2016. All patients signed informed consent, and samples were acquired under the University of Pittsburgh IRB approved protocol (IRB0502025). We previously reported on the detection of ESR1 mutations in cfDNA from these 7 patients using ddPCR [11]. Serial blood draws (range; 2-5) were available for 4 patients. A total of 14 blood samples from 7 patients were utilized for cfDNA and buffy coat DNA isolation, followed by NGS testing.

### Patient Sample Processing

CfDNA was isolated as described previously [11]. Blood was processed to separate plasma and buffy coat by double centrifugation within 4 days of blood collection. 1ml to 4ml of plasma was used for isolation of cfDNA using QIAamp Circulating Nucleic Acid kit (Qiagen). cfDNA was quantified using Qubit dsDNA HS assay kit (ThermoFisher Scientific). Genomic DNA was isolated from buffy coat using DNeasy Blood & Tissue Kit (Qiagen) for use as germline DNA control. Buffy coat DNA was quantified using Qubit dsDNA BR assay kit (ThermoFisher Scientific).

### Ion Torrent Sequencing

20ng of DNA (10ng per amplicon pool) was used for library preparation using Ion AmpliSeq™ Library Kit 2.0 (Thermo Fisher Scientific) and the custom designed MammaSeq™ primer panel (Supplementary Data File 1). Template preparation by emulsion PCR and enrichment was performed on the Ion OneTouch 2 system (ThermoFisher Scientific). Template positive Ion Sphere particles (ISP) were loaded onto Ion chips and sequenced. Tumor DNA and cfDNA samples were sequenced using P1 chips (60 million reads) on the Ion Proton™ (ThermoFisher Scientific) at empirical depths of 1000x and 5000x respectively. Buffy coat DNA was sequenced using 318 chip (6 million reads) on the Ion Torrent Personal Genome Machine (PGM™) (ThermoFisher Scientific) at 500x.

### Variant Calling

Ion Torrent Suite V4.0 was used to align raw fastq files to the hg19 reference genome and generate VCF files (4.0% AF cutoff for tumor samples, 1.0% AF cutoff for cfDNA samples). Cravat CHASM-v4.3 (http://hg19.cravat.us/CRAVAT/) was used to annotate variants with resulting protein changes and snp annotation from ExAC [12] and 1000Genomes [13]. Variant calls from buffy coat DNA were used to remove germline variants from the 14 cfDNA samples in a patient matched manner. SNP and sequencing artifact filtering, data organization, and figure preparation were performed in R (v3.4.2). The R package ComplexHeatmaps was used to generate figures 1 and 3A [14]. CNVKit was used to call copy number across all genes, however only genes containing more than 3 amplicons were reported (Table 2) [15]. DNA from the buffy coat of the cfDNA cohort was used to generate a single copy-number reference which was used as a baseline for copy number calling on the solid tumor cohort. CNKit reports copy number as a log2 ratio change. CNV were considered significant above an absolute copy number above 6 (log2(6/2)=1.58) or below 1 (log2(1/2) = −1).

**Fig 1:**
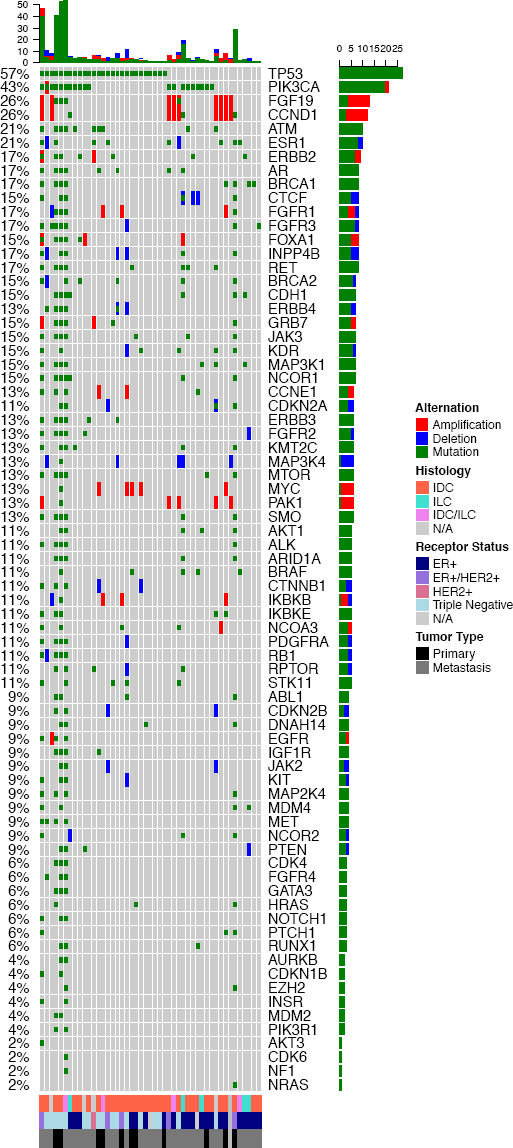
Genetic alterations identified by the MammaSeq™ gene panel in a test cohort of 46 breast cancers. Oncoprint depicting the distribution of somatic mutations, copy-number amplifications (absolute copy-number greater than 6), and deletions (absolute copy-number less than 1).

**Table 2:** 79 Genes incorporated in the MammaSeqTM gene panel.

|  |  |  |  |  |
| --- | --- | --- | --- | --- |
| ABL1 | CDK6 | FGFR3 | KDR | NOTCH1 |
| AKT1 | CDKN1B | FGFR4 | KIT | NRAS |
| AKT3 | CDKN2A | FOXA1 | KMT2C | PAK1 |
| ALK | CDKN2B | GATA3 | KRAS* | PDGFRA |
| AR | CTCF | GRB7 | MAP2K4 | PIK3CA |
| ARID1A | CTNNB1 | HIST2H2BE* | MAP3K1 | PIK3R1 |
| ATM | DNAH14 | HRAS* | MAP3K4 | PTCH1 |
| AURKA | EGFR | IDH1* | MDM2 | PTEN |
| AURKB | ERBB2 | IGF1R | MDM4 | RB1 |
| BRAF | ERBB3 | IKBKB | MET | RET |
| BRCA1 | ERBB4 | IKBKE | MTOR | RPTOR |
| BRCA2 | ESR1 | INPP4B | MYC | RUNX1 |
| CCND1 | EZH2* | INSR | NCOA3 | SMO |
| CCNE1 | FGF19 | JAK2 | NCOR1 | STK11 |
| CDH1 | FGFR1 | JAK3 | NCOR2 | TP53 |
| CDK4 | FGFR2 | JUN* | NF1 |  |

\* denotes genes with less than 3 amplicons, for which copy number changes were not reported

### Data and code

Annotated, unfiltered, mutation and CNV data, along with R code related to this study, are deposited on GitHub. (https://github.com/smithng1215)

### ddPCR

2 ng of cfDNA or buffy coat DNA was subjected to targeted high-fidelity preamplification for 15 cycles using custom designed primers (Supplemental Table 2) and PCR conditions previously described [11]. Targeted preamplification products were purified using QIAquick PCR Purification kit (Qiagen) and diluted at 1:20 before use in ddPCR reaction. 1.5ul of diluted preamplified DNA was used as input for ddPCR reaction. ddPCR was performed for ESR1-D538G, FOXA1-Y175C, and PIK3CA-H1047R mutations. Custom ddPCR assays were developed for ESR1-D538G (Integrated DNA Technologies) and FOXA1-Y175C (ThermoFisher Scientific). Sequences are described in Supplementary Table 3. PIK3CA-H1047R was analyzed using PrimePCR ddPCR assay (Bio-Rad Laboratories) dHsaCP2000078 (PIK3CA)/ dHsaCP2000077 (H1047R). Nuclease-free water and buffy coat-derived wildtype genomic DNA as negative controls, and oligonucleotides carrying mutation of interest or DNA from a cell line with mutation as positive controls were included in each run to eliminate potential false positive mutant signals. An allele frequency of 0.1% was used as a lower limit of detection.

### Statistical Analysis

All statistical analysis was performed in R 3.4.2. To determine if there was a significant correlation between mutational burden and copy number burden, we calculated the pearson correlation coefficient between the number of somatic mutations in each sample, with the number of significant copy number changes in each sample.

## Results

### Development of MammaSeq™ Panel

To build a comprehensive list of somatic mutations in breast cancer, we combined mutation calls from primary tumors in TCGA (curated list level 2.1.0.0) and limited studies focused on metastatic breast cancer [16-18]. The biological function and druggablity of mutated genes were investigated via Gene Ontology (GO) [19] and DGIdb (v2.0) databases [20]. The information regarding FDA approved drugs was downloaded from “https://www.fda.gov/Drugs” and added to our list. We used the following criteria to priotrize the clinically important mutated genes:

- The mutated gene is among significantly mutated genes (SMGs) in primary and metastatic samples.
- The mutated gene is clinically actionable (e.g. there is available FDA-approved drug(s) against it).
- The mutated gene is of functional importance in cancer (e.g. kinase genes were scored higher in the list).
- The mutation has been found in more than 5 primary tumors OR 2 metastatic tumors.
- The mutation has been found in both primary and metastatic lesions.

The final mutation list was then curated and narrowed down to 80 genes and 1398 mutations. Additional amplicons were added to select genes to ensure sufficient coverage of genes known to harbor functional copy-number variants. Amplicon probe design was unsuccessful for 29 mutations, including all 3 mutations in the gene HLA-A, yielding a final panel consisting of 688 amplicons targeting 1369 mutations across 79 genes. (Selected genes described in Table 2. Gene coverage depicted in Supplemental Figure 1. Panel design described in supplemental data file 1).

The panel includes 34 of the 50 (68%) genes incorporated in AmpliSeq Cancer Hotspot Panel v2. Genes that were not mutated in breast cancer (TCGA and in-house data) and genes that were not considered to be clinically actionable were not included. The MammaSeq™ panel includes 8 of the 10 (80%) genes and ~ 91% of the hotspots targeted by the Thermo Oncomine Breast cfDNA assay. MammaSeq™ covers 14% of the base pairs covered by the Qiagen Human Breast Cancer GeneRead DNAseq Targeted Array, however, it covers hotspots in over half of the genes (57%) (plus an additional 34 genes). Of these panels, MammaSeq is the only one that includes CDK4 and CDK6, both of which can be targeted with FDA approved CDK4/6 inhibitors [21]. Additional genes unique to MammaSeq include common drivers, CCND1, MTOR, and FGFR4. Finaly, MammaSeq covers 68 of 315 genes targeted by the larger pan cancer Foundation Medicine, FoundationOne panel. Supplemental figure 2 details the overlap in coverage between MammaSeq™ and above mentioned commercially available panels.

### Characterization of Genetic Variants detected by Mammaseq in a Solid Tumor Cohort

To evaluate performance in mutation detection by the MammaSeq™ panel, sequencing was carried out on a cohort of 46 solid tumor samples, with a mean read depth of 2311X (Supplemental Figure 3). 4970 total variants (mean: 106, median: 82) were called across all patient samples. We removed identical genomic variants that were present in more than 10 samples as these were likely to be sequencing artifacts or common SNPs. Removing non-coding and synonymous variants yielded 1433 and 901 variants, respectively. To filter out less common polymorphisms, we removed variants annotated in ExAC [12] or the 1000Genomes [13] databases in more than 1% of the population. We removed variants with an allele frequency above 90% as these were likely germline. Finally, to focus on high confidence mutations, we removed variants with a strand bias outside of the range of 0.5-0.6, yielding a total of 592 protein coding mutations (mean 12.9, median 3, IQR 3) (Figure 1).

Interestingly, as noted by the variation between the mean and median, the total number of mutations was skewed toward a subset of samples (Figure 1-top panel). 408 of the 592 mutations (69%) were found in just 4 of the 46 samples (Supplemental Figure 4). These 4 samples are by definition outliers, as they are all more than 1.5 times the IQR plus the median. 3 of these 4 samples with high mutational burden were of triple negative subtype, the fourth being ER+/HER2+. The most common mutated genes were TP53 (57%) and PIK3CA (43%). We also noted common mutations in ESR1 (21%), ATM (21%) and ERBB2 (17%).

To examine CNV changes, we established a baseline for pull down and amplification efficiency by performing MammaSeq™ on normal germline DNA from 14 samples (7 patients − 6 additional). CNVkit [15] was used to pool the normal samples into single reference and then call CNV in the solid tumor cohort (Figure 1). CNV were identified in many common oncogenes including *CCND1, MYC, FGFR1* and others. 2 of the 3 *ERBB2^+^* samples (via clinical assay) showed CNV by MammaSeq. FGF19 and CCND1 were co-amplified in 9 of the 46 (20%) solid tumors. Both genes are located on 11q13, a band identified in GWA studies as harboring variants, including amplifications, associated with ER+ breast cancers [22]. There wasn’t a correlation between mutational burden and copy number burden (pearson correlation p-value = 0.7445).

### Clinical Utility of Genetic Variants Detected by MammaSeq

To determine how many of the mutations have putative clinical utility, we utilized the OncoKB precision oncology knowledge database [23]. 25 of the genes in the MammaSeq™ panel (32% of the panel) harbor clinically actionable variants with supporting clinical evidence (OncoKB levels 1-3). In total, we identified 28 actionable variants (26 SNV and 2 ERBB2 amplifications) that have supporting clinical evidence (level 1-3) and an additional 3 actionable variants supported by substantial research evidence (level 4) in the solid tumor cohort (Table 3). The 26 SNVs were distributed across 20 of the 46 cases (43%) (Figure 2). Consistent with the report detailing the development of the OncoKB database [24], the vast majority of actionable variants in breast cancer are annotated at level 3, indicating that variants have been used as biomarkers in Clinical Trials, however they are not FDA approved. In fact, the only level 1 annotated variant in breast cancer is *ERBB2* amplification.

**Fig 2:**
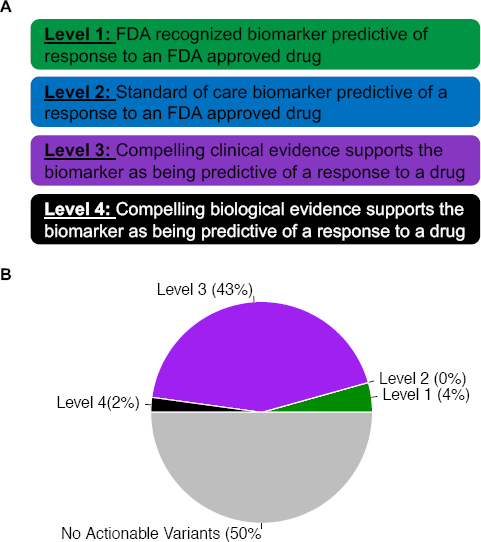
Clinical Actionality of MammaSeq™ identified somatic alterations. **(A.)** Annotation levels, adapted from OncoKB[23] **(B.)** Samples were categorized based on the most actionable alteration. Specific alterations and associated drugs are depicted in Table 3.

**Table 3:** Identified variants in annotated in OncoKB with corresponding targeted therapeutics.

| Sample ID | Gene | Protein Sequence Change | Allele Frequency | Level | Drugs |
| --- | --- | --- | --- | --- | --- |
| MET_03 | ERBB2 | Amplification | - | 1 | Lapatinib + Trastuzumab, Pertuzumab + Trastuzumab, Ado-trastuzumab emtansine, Lapatinib, Trastuzumab |
| MET_33 | ERBB2 | Amplification | - | 1 |  |
| MET_39 | AKT1 | E17K | 0.25 | 3 | AZD5363 |
| MET_18 | ERBB2 | I654V | 0.122222 | 3 | Neratinib |
| MET_32 | ERBB2 | I654V | 0.461731 | 3 |  |
| MET_49 | ERBB2 | I654V | 0.495495 | 3 |  |
| MET_07 | ESR1 | D538G | 0.477717 | 3 | AZD9496, Fulvestrant |
| MET_21 | ESR1 | D538G | 0.335884 | 3 |  |
| MET_28 | ESR1 | D538G | 0.454271 | 3 |  |
| MET_27 | ESR1 | Y537S | 0.376441 | 3 |  |
| MET_22 | PIK3CA | E453K | 0.444722 | 3 | Buparlisib, Serabelisib, Alpelisib + Fulvestrant, Copanlisib, GDC-0077, Alpelisib, Taselisib + Fulvestrant, Buparlisib + Fulvestrant, Taselisib |
| MET_10 | PIK3CA | E542K | 0.106212 | 3 |  |
| MET_21 | PIK3CA | E542K | 0.501912 | 3 |  |
| MET_41 | PIK3CA | E542K | 0.073183 | 3 |  |
| MET_49 | PIK3CA | E542K | 0.467702 | 3 |  |
| MET_08 | PIK3CA | E545K | 0.204327 | 3 |  |
| MET_34 | PIK3CA | E545K | 0.0871914 | 3 |  |
| MET_40 | PIK3CA | E545K | 0.844344 | 3 |  |
| MET_25 | PIK3CA | H1047R | 0.341171 | 3 |  |
| MET_29 | PIK3CA | H1047R | 0.180681 | 3 |  |
| MET_32 | PIK3CA | H1047R | 0.2785 | 3 |  |
| MET_33 | PIK3CA | H1047R | 0.413998 | 3 |  |
| MET_38 | PIK3CA | H1047R | 0.384692 | 3 |  |
| MET_44 | PIK3CA | H1047R | 0.60054 | 3 |  |
| MET_06 | PIK3CA | N345K | 0.376571 | 3 |  |
| MET_35 | PIK3CA | Q546R | 0.435484 | 3 |  |
| PR_26 | BRAF | G469A | 0.52028 | 4 | LTT462, BVD-523, KO-994 |
| MET_34 | KRAS | G12D | 0.074 | 4 | LY3214996, KO-947, GDC-1014 |
| MET_22 | PTEN | C136Y | 0.756233 | 4 | AZD6482 + Alpelisib |
| MET_01 | PTEN | R130Q | 0.116279 | 4 |  |
| CF_28_Draw_1 | ESR1 | D538G | 0.0746562 | 3 | AZD9496, Fulvestrant |
| CF_28_Draw_5 | ESR1 | D538G | 0.146853 | 3 |  |
| CF_22_Draw_1 | PIK3CA | H1047R | 0.320088 | 3 | Buparlisib, Serabelisib, Alpelisib + Fulvestrant, Copanlisib, GDC-0077, Alpelisib, Taselisib + Fulvestrant, Buparlisib + Fulvestrant, Taselisib |
| CF_22_Draw_2 | PIK3CA | H1047R | 0.402402 | 3 |  |

### Characterization of Genetic Variants detected by Mammaseq in cfDNA

To examine the potential of MammaSeq™ to detect variants in cfDNA, we sequenced 14 cfDNA samples isolated from 7 patients with metastatic disease. cfDNA samples were sequenced to a mean depth of 1810X, while matched buffy gDNA was sequenced to a mean depth of 425X (Supplemental figure 4).

We applied the same filtering pipeline to the cfDNA variants and solid tumor variants, except in the smaller cohort we removed all identical variants found in more than 4 samples, and lowered the minimum allele frequency to 1.0%. We identified a total of 43 somatic mutations across the 14 cfDNA samples (mean: 3.1, median 1, IQR 1.75) (Figure 3A). Similar to the solid tumor cohort, a single draw from 1 patient (CF_28- Draw 1) harbored 25 of the 13 (58%) total mutations. Using the same definition, this sample is also an outlier. Similar to the solid tumor cohort, PIK3CA and ESR1 were among the most commonly mutated genes.

**Fig 3:**
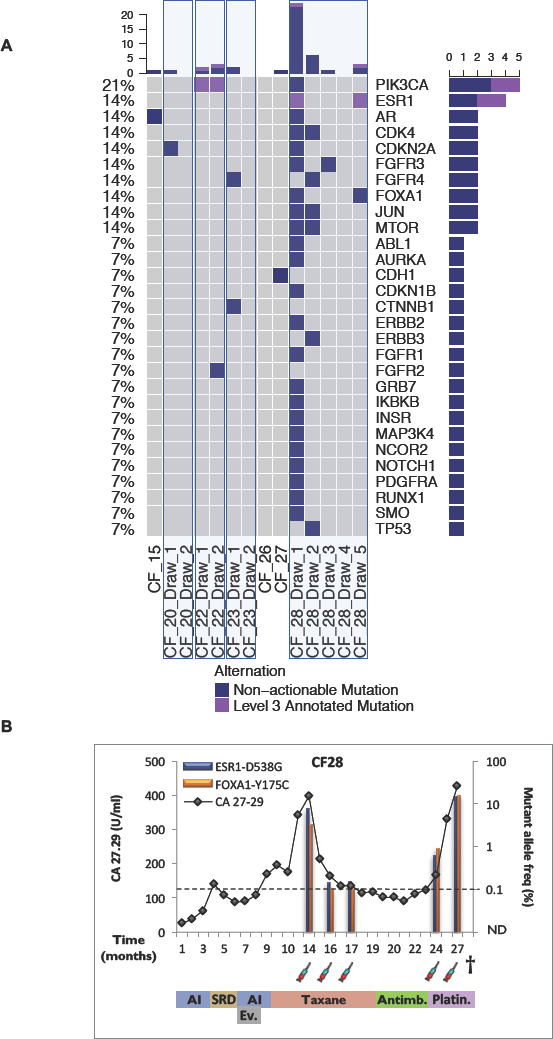
Genetic alterations identified in cfDNA from a test cohort of 7 patients with metastatic invasive ductal carcinoma. **(A.)** Oncoprint of somatic mutations identified in 14 cfDNA samples. **(B.)** Clinical timeline and mutant allele frequency of ESR1-D538G and FOXA1-Y175C mutations in serial blood draws from patient CF28. The timeline starts with diagnosis of metastasis and shows tumor marker assessments (CA 27.29 antigen line graph), mutant allele frequency (bar graphs), LLoD (dotted line), blood draws (syringe), and treatments received. Treatment abbreviations: AI (aromatase inhibitor), SERD (selective estrogen receptor degrader), Ev. (Everolimus), Antimb. (Antimetabolite), Platin (Platinum-based chemotherapy).

Two of the identified somatic mutations (each identified in 2 draws from 1 patient) are annotated at level 3 in the OncoKB database, ESR1 - D538G and PIK3CA - H1047R (Figure 3A). The ESR1 mutation was identified in 2 separate blood draws from patient CF_28 taken 13 months apart. Interestingly, the FOXA1 - Y175C mutation was also identified in the same draws from patient CF_28 (Figure 3B). The allele frequencies of these mutations strongly correlate with levels of cancer antigen 27-29 (CA-27.29), indicating that the mutation frequencies are likely an indicator of disease burden. Mutations identified in all three genes (ESR1, PIK3CA, and FOXA1) were independently validated using ddPCR (Supplemental Figure 5).

## Discussion

Advances in the accuracy, cost, and analysis of NGS make it an ideal platform to develop diagnostics that can be used to precisely identify treatment options. MammaSeq was developed to comprehensively cover known driver mutation hotspots specifically in primary and metastasis breast cancer that would identify mutations with potential prognostic value. Typically, NGS diagnostics are reserved for late stage disease. As a result, (as noted in our previous publication[8]), the solid tumor cohort was significantly enriched for metastatic disease and markers of poor prognosis - triple negative subtype, late presentation, and therapy resistance.

Consistent with previous mutational studies, we report that a small subset of breast cancers harbor high mutational burden [25]. Across a variety of cancers, groups have demonstrated the correlation between the tumor mutation burden (TMB) and the efficacy of immunotherapy checkpoint inhibitors (reviewed here [26]). However, the ability to accurately depict tumor mutation burden is dependent on the percentage of the covered exome. Illumina have shown that the TruSight Tumor 170 panel (170 genes and 0.524 Mb) begins to skew the TMB upwards, when used on samples that contain relatively few mutations [27]. A previous study by Chalmers et al. used a computational model to show that below 0.5Mb, TMB measurements are highly variable and unreliable [28]. The MammaSeq™ panel covers just 82,035bp (0.08Mb), and we speculate that it cannot be used to calculate a mutational burden comparable to whole exome based studies. That being said, the stark difference in the total number of mutations identified in the subset of 4 tumor samples, suggests that they may be suited for immunotherapy.

Liquid biopsies are beginning to be utilized clinically after numerous proof-of-principle studies have demonstrated the potential of circulating cell-free DNA (cfDNA) for prognostication, molecular profiling, and monitoring disease burden [11, 29-33]. We have demonstrated that the MammaSeq™ panel can be used to identify mutations in cfDNA. For one patient (CF_28), we have cfDNA data from 5 blood draws taken over the course of 13 months. The sharp drop-off in the number of somatic mutations identified between the first and second draws co-occurs with a decrease in CA.27.29 levels, suggesting that the patient may have responded well to treatment, leading to disappearance of sensitive clones. In the later blood draws, we did not observe an increase in the total number of somatic mutations, however, we did find an increase in the allele frequency of ESR1-D538G and FOXA1-Y175C mutations, which may be caused by therapeutic selection of resistant clones.

High-throughput genotyping of solid tumors and continual monitoring of disease burden through sequencing of cfDNA represent potential clinical applications for NGS technologies. It should be noted that targeted DNA sequencing panels such as MammaSeq™ are far less comprehensive than whole exome sequencing and they do not allow for evaluation of structural variants, which can often lead to gene fusions that function as drivers [34]. Nevertheless, as a focused panels represent cost-effective and useful alternatives to whole exome sequencing for targeted mutation detection.

## Conclusions

Here we report the development of MammaSeq™, a targeted sequencing panel designed based on current knowledge of the most common, impactful, and targetable drivers of metastatic breast cancer. This data provides further evidence for the use of NGS diagnotsics in the management of advanced breast cancers.

## List of Abbreviations

CfDNA: circulating-free cfDNA
CNV: Copy Number Variants
DdPCR: Droplet Digital PCR
DgIDB: Drug-Gene Interaction Database
GDNA: Genomic DNA
GO: Gene Ontology
GWAs: Genome Wide Association studies
IDC: Invasive Ductal Carcinoma
ILC: Invasive Lobular Carcinoma
ISP: Ion Sphere particles
MBC: Metastatic Breast Cancer
NGS: Next Generation Sequencing
PGM: Ion Torrent Personal Genome Machine
SMG: Significantly Mutated Gene
SNP: Single Nucleotide Polymorphism
SNV: Single Nucleotide Variant
TCGA: The Cancer Genome Atlas
TMB: Tumor Mutational Burden

## Declarations

### Ethics approval and consent to participate

The research was performed under the University of Pittsburgh IRB approved protocol PRO16030066.

### Consent for publication

Not applicable.

### Availability of data and material

Annotated, unfiltered, mutation and CNV data, along with R code related to this study, are deposited on GitHub (https://github.com/smithng1215).

### Competing Interests

RJH received salary and has ownership interest (including patents) in Foundation Medicine and is currently an employee at AstraZeneca. Other authors declare that they have no conflict of interests to report.

### Funding

Research funding for this project was provided in part by a Susan G. Komen Scholar award to AVL and to SO, the Breast Cancer Research Foundation (AVL and SO), the Fashion Footwear Association of New York (SO and AVL), and a research grant from Glimmer of Hope.

### Author Contributions

RJH, AB and AVL designed the MammaSeq panel. NGS and RG analyzed data and wrote manuscript. AMB, SP, and KK collected samples. AIW, PCL, and GG performed sample processing, quality control, and sequencing. NGS, RG, and AVL analyzed data and wrote the manuscript. SO, YEN, and MNN provided critical feedback on panel design and manuscript writing. All authors read and approved the final manuscript.

## Acknowledgements

We are thankful to the patients who generously provided tumor tissue for our studies and to the surgical, pathology, and tissue bank colleagues for their substantial assistance and support. This project used the University of Pittsburgh HSCRF Genomics Research Core, and the UPCI Tissue and Research Pathology Services that is supported in part by award P30CA047904. We thank Lousie Mazur and the UPMC Cancer Registry for clinical abstraction.

## Figure Legends

**Supplemental Figure 1:** MammaSeq™ gene coverage. The percentage of protein coding bases pairs in each gene that is sequenced by the MammaSeq™ panel.

**Supplemental Figure 2:** Coverage overlap between MammaSeq™ and select commercially available panels used in breast cancer. Overlap of genes present in the MammaSeq™ panel and the **(A.)** Foundation Medicine FoundationOne panel **(B.)** Thermo Ion AmpliSeq Cancer Hotspot Panel (v2) **(C.)** Qiagen GeneRead Human Breast Cancer Panel and the **(D.)** Thermo Oncomine Breast cfDNA Assay. Overlap of the number of base pairs covered for the **(E.)** Qiagen GeneRead and **(F.)** Thermo Oncomine panels were calculated as these panel designs are publicly available.

**Supplemental Figure 3:** Mean sequencing read depth for **(A.)** the 46 solid tumor cohort. **(B.)** isolated mononuclear cells from the 14 cfDNA draws and **(C.)** the 14 cfDNA samples.

**Supplemental Figure 4:** Tumor mutational burden across all samples in the 46 solid tumor cohort. **(A.)** Total detected mutations for each sample.

**Supplemental Figure 5:** ddPCR validation of mutations identified by MammaSeq™ is indicated along with mutant allele frequencies for **(A.)** ESR1-D538G, **(B.)** FOXA1- Y175C, and **(C.)** PIK3CA-H1047R.

